# Targeting IL-2 to inflamed tissues via oxidation-specific epitopes enables third-generation bispecific IL-2 therapeutics

**DOI:** 10.64898/2026.09.10.750022

**Authors:** Thomas Vazquez, Monica Cuevas-Martinez, Nicolas Billiald, Florentina Porsch, Marie Sion, Fatima Saddouk, Céline Albalaa, Lorena Zentilin, Lauri Peil, Joan Teyra, Tahar Bouceba, Ines Leleu, Sylviane Pied, Mart Ustav, Alain Tedgui, Serena Zacchigna, Roger Le Grand, Michelle Rosenzwajg, Ziad Mallat, Christoph J. Binder, David Klatzmann

## Abstract

Interleukin-2 (IL-2) is essential for the survival and activation of regulatory T cells (Tregs). Low-dose native IL-2 (IL-2LD) therapy restores immune regulation in vivo and has shown reproducible clinical benefit across multiple autoimmune, inflammatory, and neuroimmune diseases. Attempts to improve IL-2 through engineered variants (muteins) have mainly focused on enhancing Treg selectivity by reducing IL-2 receptor β-chain binding, but this strategy profoundly diminishes biological potency, likely contributing to the limited clinical efficacy of IL-2 muteins. Here, we develop a “third-generation IL-2” that combines site-specific targeting and bifunctionality. We generated a bivalent fusion protein linking IL-2 to a single-chain antibody recognizing oxidation-specific epitopes (OSEs), which are abundantly expressed at inflamed sites. Targeting OSEs provides not only site-specific localization, but also true bifunctionality as both anti-OSE antibodies and IL-2_LD_ independently show therapeutic benefit in limiting inflammation. We first show that IL-2_IT_ has bifunctional biological activities in vitro. In vivo, IL-2_IT_ had increased specificity for Treg over Teff activation, which we attribute to a conformation-dependent modulation of IL-2 receptor engagement. Importantly, IL-2_IT_ provided precise delivery to inflamed tissues in models of psoriasis and colitis. Altogether, this resulted in superior therapeutic benefit in multiple clinical settings, including in atherosclerosis models. Thus, our strategy illustrates a generalizable approach to cytokine engineering that preserves native signaling while achieving spatial control. Specifically, our findings validate OSE targeting as an efficient strategy to guide therapeutics to sites of inflammation and establish OSE-IL-2 as a promising bispecific Treg engager for treating inflammation.

**One sentence summary:** Targeted delivery of IL-2 to inflamed sites via oxidation-specific epitope recognition establishes a third generation of bispecific IL-2 therapeutics.

## Introduction

Interleukin-2 (IL-2) is a pleiotropic cytokine with a non-redundant role in immune homeostasis(*1–3*). At physiological concentrations, IL-2 is essential for the survival, lineage stability, and suppressive function of regulatory T cells (Tregs)(*4*), which restrain excessive immune activation and maintain self-tolerance(*5–7*). Therapeutic exploitation of this biology through low-dose IL-2 (IL-2_LD_) administration has revealed a unique capacity to preferentially activate and expand Tregs in vivo. At doses up to 3 million international units (MIU), IL-2_LD_ induces little to no activation of effector T cells (Teffs)(*8*), thereby shifting the immune balance toward regulation. Over the past decade, IL-2_LD_ has demonstrated consistent immunological activity and signals of clinical benefit across multiple autoimmune, inflammatory, and neuroimmune diseases, including systemic lupus erythematosus(*9–12*), type 1 diabetes(*13, 14*), graft-versus-host disease(*15, 16*), amyotrophic lateral sclerosis(*17*), and atherosclerotic cardiovascular disease(*18, 19*), establishing the potential of native IL-2 as a Treg-stimulatory therapy in humans.

Motivated by these advances, extensive efforts have focused on engineering IL-2 variants with improved selectivity and pharmacokinetic properties. Most IL-2 muteins have been designed to reduce affinity for the IL-2 receptor β chain (CD122), thereby biasing signaling toward the high-affinity trimeric IL-2 receptor constitutively expressed by Tregs(*20–22*). However, this strategy intrinsically compromises IL-2 signaling strength, often by 100- to a 1000-fold when assessed by pSTAT5 phosphorylation. In clinical settings, such attenuation has translated into limited therapeutic efficacy, despite numerical expansion of Tregs. This phenomenon has been particularly evident in systemic lupus erythematosus (SLE), where multiple phase 2 trials of native IL-2_LD_ have reported positive immunological and clinical outcomes, leading to regulatory approval and marketing of IL-2_LD_ therapy in China based on national regulatory criteria. In contrast, programs deploying engineered IL-2 variants have so far failed to yield efficacy in SLE. Although no results have been published yet, Celgene, Moderna, Lilly, and Amgen (https://www.fiercebiotech.com/biotech/amgen-discontinues-rd-work-lupus-futility-reasons) have all announced the termination of their program in SLE. These outcomes expose a central constraint of current IL-2 engineering paradigms, at least when based primarily on receptor-affinity attenuation. Collectively, these findings suggest that the primary challenge in IL-2 therapeutics is not excessive pleiotropy, but insufficient control over the spatial and contextual delivery of intact IL-2 signaling.

Accordingly, further progress in IL-2–based immunotherapy is unlikely to arise from additional reductions in receptor affinity. Instead, next-generation IL-2 therapeutics should preserve native cytokine potency while incorporating new functional properties, such as tissue targeting, cell-type restriction, or bifunctionality. By directing IL-2 activity to defined pathological niches—rather than globally dampening its signaling—such approaches offer the prospect of enhancing efficacy and precision without sacrificing the favorable safety profile established by IL-2_LD_.

Oxidation-specific epitopes (OSEs) provide an attractive molecular cue for such spatial targeting. OSEs are generated during oxidative stress and tissue injury and are abundantly exposed at sites of chronic inflammation and autoimmune pathology across species(*23*). As damage-associated molecular patterns, OSEs are recognized by natural antibodies and scavenger receptors, thereby flagging inflamed or injured tissues for immune regulation and repair(*24*). Targeting IL-2 to OSEs offers a strategy to localize Treg stimulation selectively to inflamed sites, increasing local immunoregulatory activity while limiting systemic exposure. In addition, antibodies directed against OSEs possess intrinsic anti-inflammatory properties, and a single-chain variable fragment (scFv) targeting OSEs has shown efficacy in preclinical models of atherosclerosis(*25–27*) and metabolic-associated fatty liver disease(*28, 29*).

Importantly, both arms of this strategy—OSE targeting and IL-2–mediated Treg activation—have independently demonstrated therapeutic relevance in atherosclerosis, a prototypical chronic inflammatory disease characterized by abundant OSE expression on oxidized lipoproteins and within arterial plaques, and by a critical role for Tregs in restraining disease progression. Anti-OSE scFv reduces plaque burden and vascular inflammation(*25*), while IL-2_LD_ enhances Treg-dependent control of vascular immune responses(*30, 31*). OSE-targeted IL-2 constructs thus exemplify a “third-generation IL-2” concept: a rationally engineered, bifunctional immunoregulatory biologic that integrates preserved cytokine potency with spatial precision, enabling more effective and context-specific immune modulation.

## Results

### Properties of a dimeric-IL-2 protein backbone

To develop an inflammation-targeted IL-2 (IL-2_IT_), we first engineered a stable IL-2 fusion protein to serve as the molecular scaffold for subsequent targeting. This construct consists of wild-type human IL-2 fused via a flexible G4S linker to a short fragment of the C4-binding protein β chain (C4bpβ) (Figure 1A). Two cysteine residues within C4BPβ mediate disulfide-bond formation between monomers, generating a dimeric IL-2 molecule (dimIL-2) with an apparent molecular weight of 46 kDa, compared to 15 kDa for native IL-2. Importantly, dimIL-2 preserves the wild-type IL-2 sequence and does not rely on receptor-affinity–altering mutations.

**Fig. 1.**
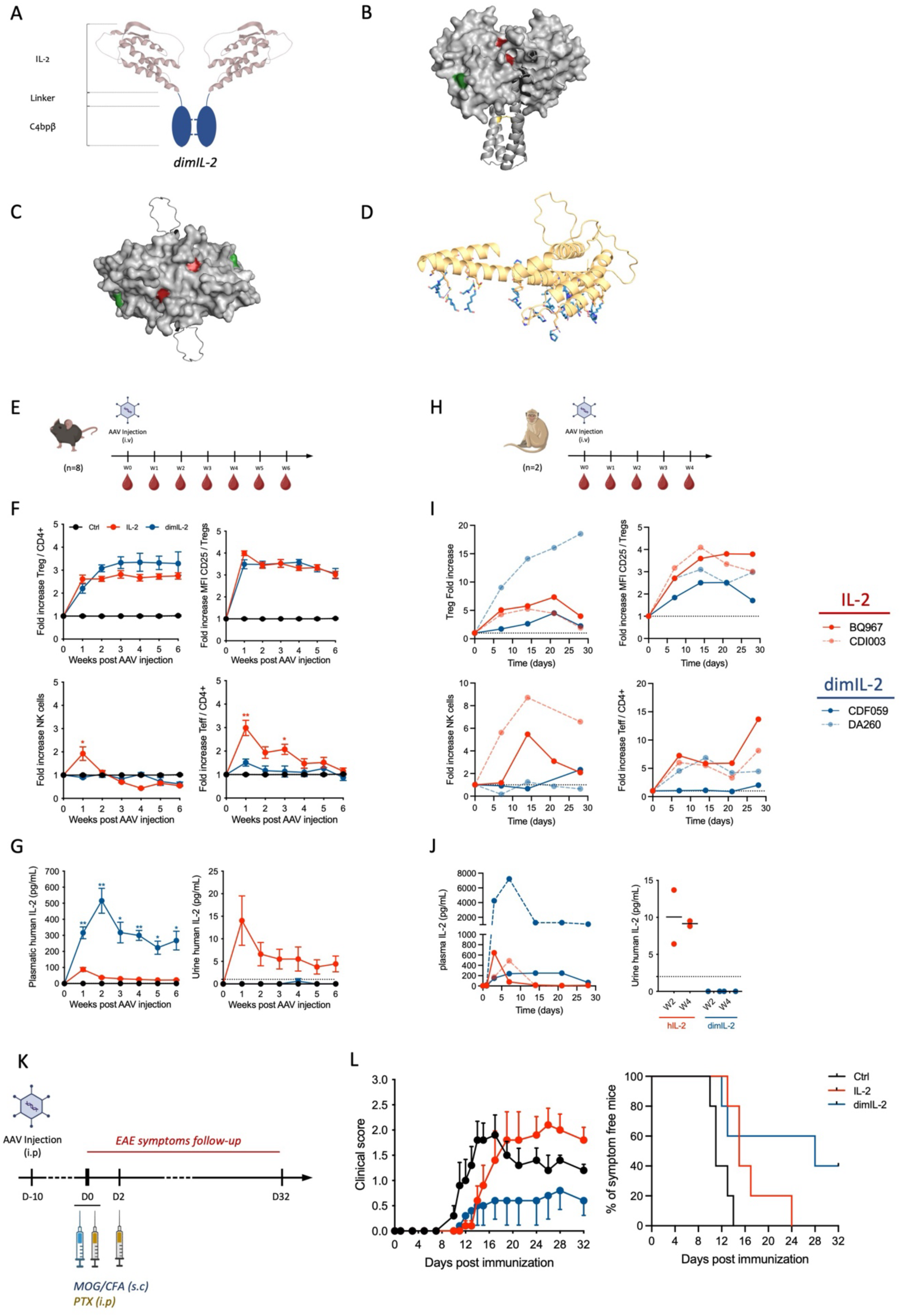
Biological properties of a dimeric IL-2 protein backbone. Schematic representation of the dimeric IL-2 (dimIL-2) protein backbone, consisting of an IL-2 molecule fused to a C4bpβ scaffold allowing dimerization **(A)**. dimIL-2 modeling using Alphafold3 indicated a conformation in which IL-2Rβ binding residues N88 and V91 (red) are enfolded while the IL-2Rα binding residue F42 (green) remains accessible, in leteral view **(B)** and in top view **(C)**. Interactions involved in the IL-2/IL-2 homodimerization are shown on one IL-2 protein at the interface **(D)**. Pharmacodynamics properties of dimIL-2 compared to native IL-2 after a single injection of 10^11^vg AAV was determined in both mice (n=8) **(E)** and non-human primate (NHP, n=2/group) **(H)**. Fold increase of Tregs, Teffs and NK cells was determined in kinetics as well as the CD25 expression among Tregs in mice **(F)** and NHP **(I)**. Pharmacokinetics was also determined in plasma (left) and urine (right), to determine urinary excretion, in both mice **(G)** and NHP **(J)**. Efficacy of dimIL-2 to prevent autoimmunity in an experimental autoimmune encephalomyelitis (EAE) was evaluated in C57BL/6J mice (n=5/group) following 10^11^vg AAV 10 days prior to MOG_35-55_ immunization **(K)**. Clinical score and percentage of symptom-free mice is plotted to determine EAE incidence and severity compared to a control group not treated **(L)**.

We used AlphaFold3 to create a structural model of dimIL-2 to evaluate the possible effects of dimerization on IL-2 receptor (IL-2Rs) binding. The predicted dimIL-2 structure shows that, due to dimerization of C4BPβ, the two IL-2 moieties may interact in a “close state” enfolding the IL-2Rβγ binding domain (Figures 1B-C). This interaction is predicted to mainly result from hydrophobic interactions and hydrogen bonds (Figure 1D and Figure S1A–B). Interestingly, the accuracy of the Alphafold3 prediction for dimIL-2 is strongly supported by the fact that it also predicted possible IL-2 homodimerization at the same IL-2Rβγ binding domain (Figure S1C), a prediction also supported by experimental crystallography(*32*) at high concentration (Figure S1D). Mechanistically, the 3D modelling showed that the dimIL-2 conformation enfolds the IL-2Rβ binding domains (Figure 1B-C, N88 and V91 positions shown in red), while maintaining the IL-2Rα binding domains accessible (F42 position shown in green) (Figure 1B-C). Such conformation could thus have significant effect on the dimIL-2 biological activity.

To get insights on dimIL-2 biological activity in vivo, we first harnessed adeno-associated viral (AAV) vectors that we previously used to express IL-2(*33*) and that spare the production and repeated administration of recombinant proteins. We generated AAVs encoding either IL-2 or dimIL-2 and administered a single intraperitoneal injection (10¹¹ vg) to mice (Figure 1E). Peripheral blood was analyzed weekly for six weeks. AAV-IL-2 induced a rapid Treg expansion detectable at week 1, plateauing at ≈3-fold above baseline. In contrast, AAV-dimIL-2 elicited a slightly slower and stronger response, reaching >3-fold expansion by week 3 (Figure 1F upper left). Both constructs similarly enhanced Treg activation, as reflected by a 3.5-fold increase in CD25 expression (Figure 1F upper right). However, IL-2 also transiently expanded NK cells and induced an up to 3-fold rise in effector CD4⁺Foxp3⁻CD25⁺ T cells (Teffs), whereas dimIL-2 had minimal effect on NK and Teff cells (Figure 1F lower left and right). Thus, the structural changes induced by dimerization on C4BPβ effectively maintain the activity of dimIL-2 towards Treg, while reducing it towards NK and T conv in vivo. Of note, while computational, AlphaFold3 predictions are consistent with the observed selective signaling bias in vivo.

Pharmacokinetic analyses revealed higher and more sustained plasma concentrations of dimIL-2 (≈300 pg/mL) compared with IL-2 (≈20 pg/mL) (Figure 1G left), that is likely due to urinary excretion of IL-2. Indeed, IL-2 was consistently detected in urine throughout the 6-week period, while dimIL-2 was undetectable at all time points (Figure 1G right). These findings indicate that dimerization increases molecular size sufficiently to prevent renal clearance.

Comparable results were obtained in cynomolgus macaques following a single intravenous injection of 5 × 10¹¹ vg/kg (Figure 1H). Longitudinal analyses over four weeks showed a sustained 5- to 15-fold Treg expansion in animals treated with AAV-dimIL-2, compared with ≤5-fold with AAV-IL-2 (Figure 1I upper left). Treg activation, assessed by CD25 upregulation, was maintained throughout the study (3.5-fold for IL-2 vs. 2.5-fold for dimIL-2)(Figure 1I upper right). As in mice, IL-2 expanded both Teffs and NK cells, while dimIL-2 largely spared these populations (Figure 1I lower left and right). One macaque treated with dimIL-2 exhibited Teff expansion similar to IL-2-treated animals, yet with a proportionally greater Treg increase. Plasma IL-2 levels were consistently higher and more stable with dimIL-2 (200–300 pg/mL plateau) compared with IL-2, which showed an early peak followed by rapid decline to ≈20 pg/mL (Figure 1J left). Again, IL-2 but not dimIL-2 was detected in urine (Figure 1J right).

Together, these results demonstrate that dimeric IL-2 retains full Treg-stimulating activity while minimizing effector-cell activation. The increased molecular size reduces renal clearance, resulting in higher and more sustained plasma levels and Treg/Teff and Treg/NK ratios consistently favoring immune regulation.

### dimIL-2 protects against autoimmunity in the EAE model

We next tested dimIL-2 in experimental autoimmune encephalomyelitis (EAE). Mice received AAV-IL-2 or AAV-dimIL-2 (10¹¹vg, intraperitoneally) ten days before EAE induction with MOG/CFA and pertussis toxin (Figure 1K). Control animals developed typical EAE with onset around day 10 and a peak clinical score of 2. AAV-IL-2 delayed onset but did not reduce severity. In contrast, AAV-dimIL-2 delayed onset and markedly attenuated disease, limiting the mean score to 0.5 through day 32 and protecting ≈40% of mice from symptoms (Figure 1L). Thus, dimIL-2 selectively activates Tregs in vivo and confers superior prevention of autoimmunity compared with IL-2.

### Design and characterization of an OSE-targeted IL-2 (IL-2_IT_-E06)

To localize Treg activation to inflamed sites, we fused the dimIL-2 backbone to a scFv derived from the prototypic E06 monoclonal antibody(*34*) that recognizes oxidized phosphocholine (OxPC) (Figure 2A). The resulting bispecific immunocytokine (IL-2_IT_-E06) has a molecular weight of ≈100 kDa and retains both IL-2 and E06 binding integrity (Figure 2B-C). A three-dimensional modelling of IL-2_IT_-E06 showed that addition of the scFv-E06 in C-terminal of dimIL-2 could maintain the closed conformation of the two IL-2s (Figure 2D), while enabling IL-2Rα binding with enough energy to break IL-2/IL-2 homodimerization and allow IL-2 docking onto the trimeric IL-2R (Figure 2E).

**Fig. 2.**
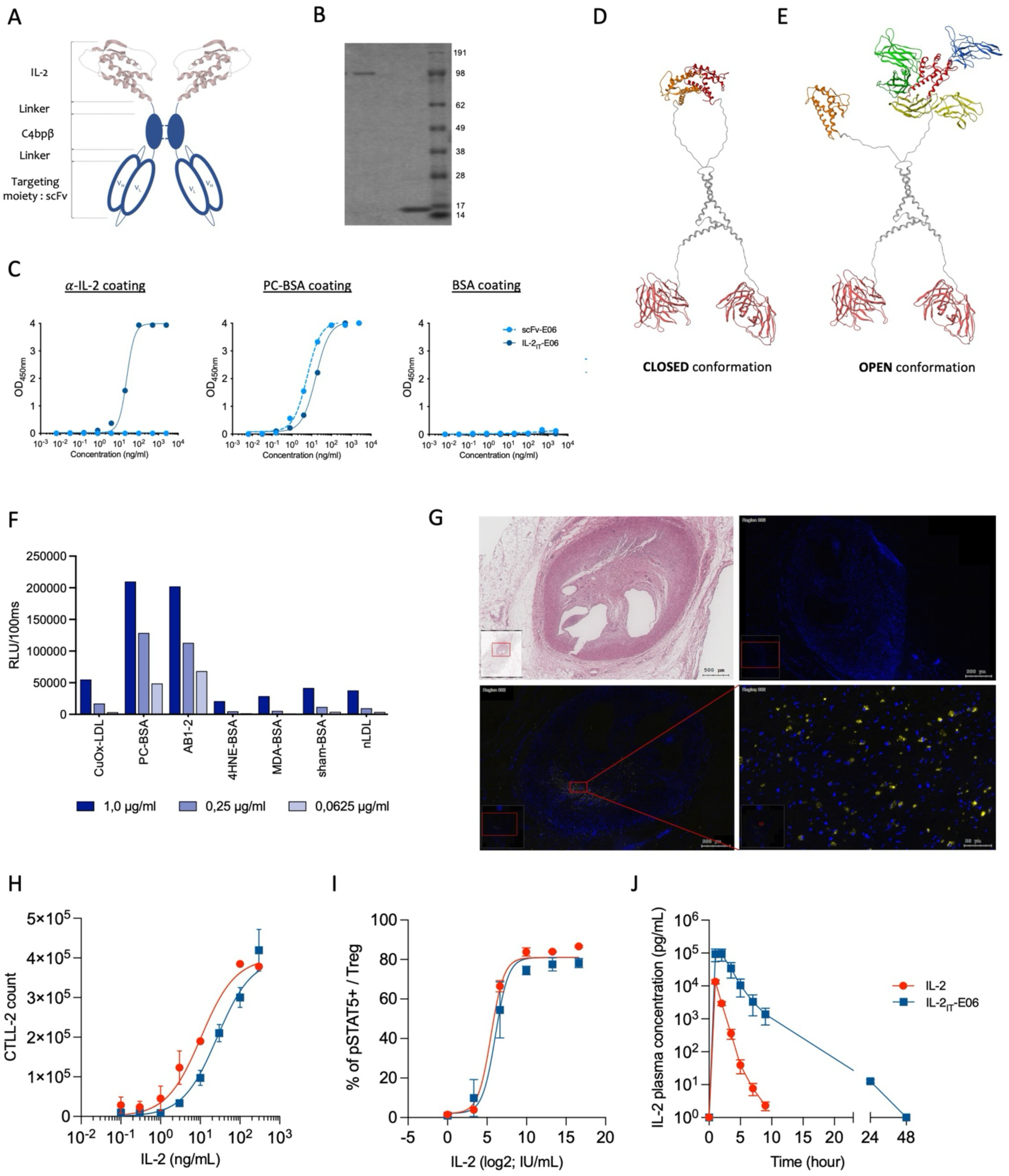
Design and characterization of an OSE-targeted IL-2 (IL-2_IT_-E06). Schematic representation IL-2_IT_-E06 consisting of the dimeric IL-2 protein backbone fused to the scFv-E06 to target OSE **(A)**. IL-2_IT_-E06 (102kDa) size increase compared to wtIL-2 (15kDa) was demonstrated in a Coomassie staining on SDS-PAGE gel **(B)**. Both scFv-E06 (dashed light blue) and IL-2_IT_-E06 (dark blue) were assessed for IL-2 detection (left), PC-BSA binding (center) or BSA binding as control (right) demonstrating IL-2_IT_-E06 is well conformed by ELISA **(C)**. AlphaFold prediction modelling illustrate the closed IL-2/IL-2 conformation in IL-2_IT_-E06 **(D)**, which could be break by the binding to the IL-2Rα, allowing a normal binding to the trimeric IL-2Rαβγ **(E)**. The binding specificity of IL-2_IT_-E06 was also evaluated against different OSEs (CuOx-LDL, idiotypic AB1-2, 4-HNE-BSA, MDA-BSA compared to Sham-BSA and native nLDL) at 1µg/ml, 0,25µg/ml and 0,0625µg/ml **(F)**. The binding capacity of IL-2_IT_-E06 was assessed in vitro on atherosclerotic cross-sections of human coronary arteries **(G)**. H&E staining (upper left) and control staining (upper right) were compared to full detection of IL-2_IT_-E06 staining (in yellow, lower left and right). Biological properties of IL-2_IT_-E06 (blue) were also compared to wtIL-2 (red) for the capacity to induce CTLL-2 cell proliferation **(H)**, induce STAT5 phosphorylation in human Tregs **(I)** and increase pharmacokinetic properties, i.e. plasmatic half-life of the protein **(J)**.

In vitro, IL-2_IT_-E06 bound selectively to OxPC bound to BSA (PC-BSA) but not to plain BSA nor to BSA coated with other class of OSE (4-HNE-BSA, MDA-BSA, or BSA) (Figure 2F). It also bounds weakly to CuOx-LDL compared to native LDL and, as expected, IL-2_IT_-E06 also bound the natural E06/T15 anti-idiotypic antibody AB1-2(*34, 35*). Importantly, in vivo, IL-2_IT_-E06 bound to both human and murine atherosclerotic plaques that are rich in OxPC(*25, 36*) (Figure 2G; Figure S2).

CTLL-2 proliferation assays showed that IL-2_IT_-E06 retained a robust IL-2 biological activity, with an IL-2 molar EC₅₀ ≈2.5-fold higher than that of IL-2 (Figure 2H). pSTAT5 phosphorylation assays also showed that IL-2_IT_-E06 retained a full potential for Treg activation (Figure 2I). In vivo, after sub-cutaneous injections, IL-2_IT_-E06 had a threefold longer half-life than IL-2 (3.7 h vs 1.1 h), with persistence beyond 24 h (Figure 2J).

### IL-2_IT_-E06 targets inflammatory sites in vivo and controls autoimmune and inflammatory diseases

We next assessed the pharmacodynamics of IL-2_IT_-E06 in vivo. C57BL6/J mice received five daily subcutaneous injections of 50,000 IU of either IL-2 or purified IL-2_IT_-E06 proteins (Figure 3A). Both treatments induced continuous and comparable Treg proliferation (up to 2.5-fold by day 5) (Figure 3B-C). CD25 expression increased more with IL-2_IT_-E06 than with IL-2 (2-fold vs. 1.5-fold) (Figure 3C), possibly reflecting the bivalent nature of the IL-2_IT_-E06. Conversely, whereas NK-cell numbers remained stable in both groups, IL-2 induced a modest activation of effector T cells, which was completely prevented by IL-2_IT_-E06 (Figure 3D).

**Fig. 3.**
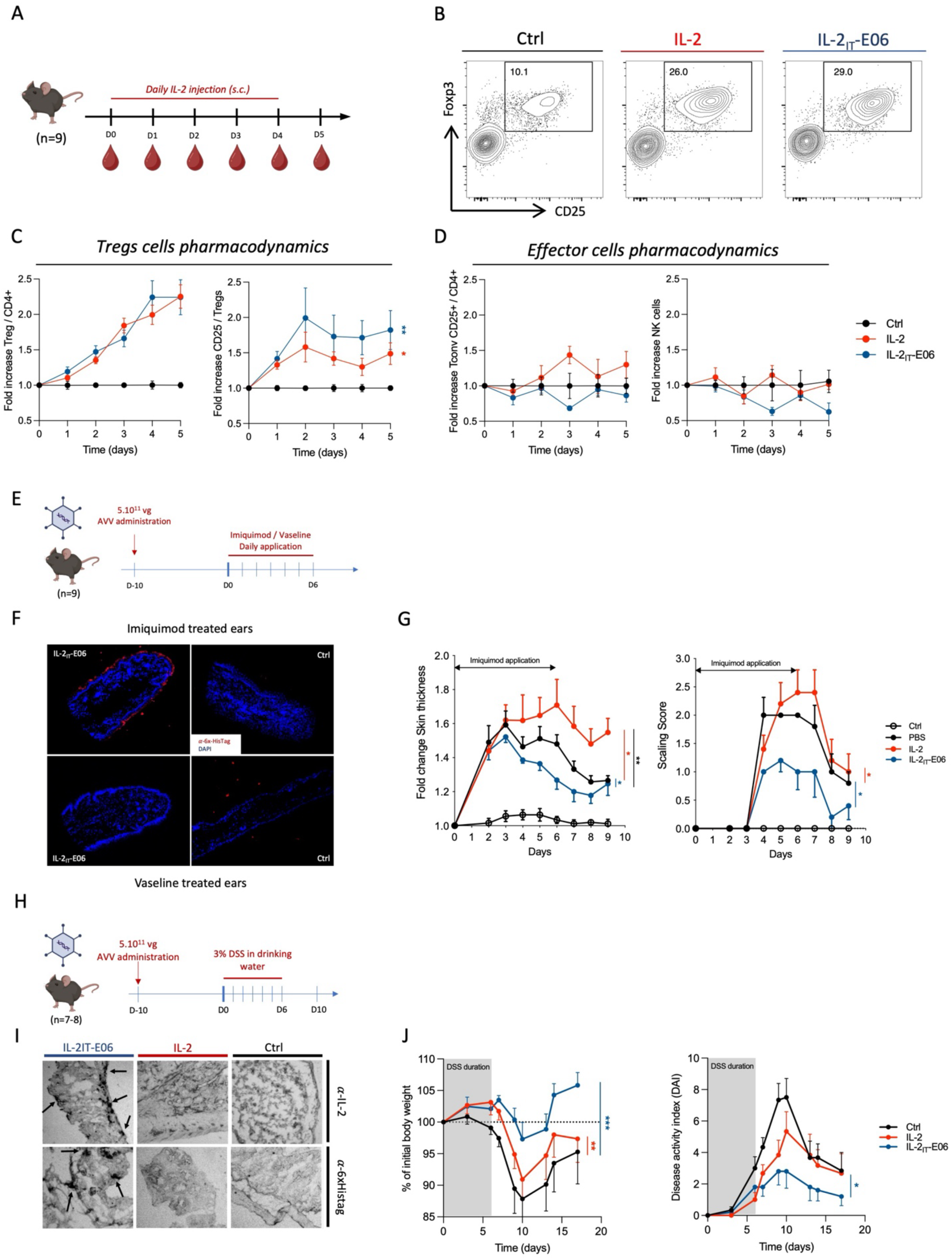
IL-2_IT_-E06 targets inflammatory sites in vivo and controls autoimmune and inflammatory diseases. IL-2_IT_-E06 recombinant protein or wtIL-2 were injected subcutaneously in C57BL/6J mice (n=9/group), 50.000 IU/day for 5 consecutive days **(A)**. Dot plots illustrating endpoint Tregs proportion among CD4+ T-cells in blood **(B)**. Blood sampling was performed 24h after each injection and pharmacodynamic changes were evaluated in Tregs **(C)** and effector cell compartments **(D)** using flow cytometry. Injection of equivalent doses in IU induced similar Treg increases for IL-2_IT_-E06 (blue) and wtIL-2 (red) **(C, left)** whereas CD25 expression is more pronounced for wtIL-2 (C, right). However, whereas NK are stable **(D, right)**, IL-2_IT_-E06 control effector T-cells expansion while wtIL-2 induced a slight increase (D, left). In vivo binding of IL-2_IT_-E06 was assessed in C57BL/6J mice (n=4/group) which have received 5% imiquimod cream application both on the shaved back and one ear, 10 days after 5.10^11^vg AAV by i.p. administration (E). After 6 days of daily imiquimod application, mice were euthanized and the two ears was sectioned and embedded in paraffin. Detection of the molecule was performed using an anti-6xHisTag antibody, co-stained with DAPI for nucleus detection **(F)**. IL-2 was only detected in the inflamed ear of mice pre-treated with AAV-IL2_IT_-E06 (F, Upper-left) while the contralateral ear of the same mice (without inflammation) did not show any staining (F, lower-left). In the same experiment, other mice (n=5/group) were followed every day for clinical symptoms as skin thickness **(G, left)** and scaling (G, right). IL-2_IT_-E06 was efficient to control psoriatic clinical symptoms while wt-IL-2 aggravated them compared to control mice without treatment. For DSS-induced colitis, C57BL/6J mice (n=7-8/group) were pre-treated with 5.10^11^vg of either AAV-IL-2_IT_-E06 or AAV-IL-2, 10 days before adding 3% DSS in drinking water for 6 days **(H)**. 2 mice per group were euthanized to evaluate in vivo binding in colon, detected in immunohistochemistry using an anti-IL-2 or an anti-6xHisTag antibody **(I)**. Other mice (n=5-6/group) were evaluated until day 17 for clinical symptoms of colitis as bodyweight loss **(J, left)** and disease activity index (DAI) (J, right).

Because the IL-2_IT_ design aims to direct IL-2 activity to inflamed tissues, we next examined its targeting and therapeutic efficacy in two distinct models of inflammation: psoriasis and colitis. In the psoriasis model, disease was induced by daily topical application of 5% imiquimod to both on the shaved back and on one ear of C57BL6/J mice, with Vaseline applied to the contralateral ear as an internal control (Figure 3E). Ten days before imiquimod treatment, mice were injected with 5.10¹¹ vg of either AAV-IL-2 or AAV-IL-2_IT_-E06; control groups included untreated mice with or without psoriasis. IL-2_IT_-E06 was visualized by immunofluorescence using an anti-6xHisTag antibody. Strong staining was detected exclusively at the border of imiquimod-treated skin, but not in Vaseline-treated control skin (Figure 3F), demonstrating that IL-2_IT_-E06 specifically localizes to inflamed sites in vivo.

Clinically, untreated mice developed typical psoriatic lesions within four days, with a 50% increase in skin thickness and a scaling score of 2. IL-2-treated mice exhibited similar early lesions but went on to develop slightly more severe disease. By day 8, they had achieved a 60% increase in thickness and a scaling score of 2.5. In striking contrast, IL-2_IT_-E06–treated mice displayed markedly reduced inflammation, with skin thickening below 25% and scaling scores ≤ 1 (Figure 3G). Thus, these findings underscore that systemic IL-2 signaling can be deleterious in certain inflammatory contexts when not spatially constrained.

We next tested IL-2_IT_-E06 in a dextran sulfate sodium (DSS)–induced colitis model. Ten days before DSS administration, mice received 5.10¹¹ vg of AAV-IL-2 or AAV-IL-2_IT_-E06 (Figure 3H); untreated mice served as positive controls for disease activity. After six days of DSS exposure, IL-2_IT_-E06 was detected in inflamed colonic tissue using both anti-IL-2 and anti-6xHisTag antibodies, whereas no staining was observed in untreated or IL-2–treated mice (Figure 3I). Functionally, IL-2 treatment provided only partial protection, delaying body-weight loss and modestly reducing disease activity (DAI ≈ 6 vs. 8 in controls). In contrast, IL-2_IT_-E06 afforded substantial protection, limiting peak weight loss to 3% and reducing the DAI to ≈ 3 (Figure 3J).

Together, these experiments demonstrate that IL-2_IT_-E06 selectively accumulates at inflammatory sites and effectively controls clinical symptoms in models of psoriasis and colitis—diseases in which conventional IL-2 either fails or worsens inflammation.

### IL-2_IT_-E06 decreases lesions size and stabilizes atherosclerotic plaques

As both IL-2 and E06-scfv showed interesting therapeutic potential in atherosclerosis(*25–27, 31, 37, 38*), we aimed to evaluate IL-2_IT_-E06 in this chronic inflammatory disease. *Ldlr⁻/⁻*mice received AAV-IL-2_IT_-E06 (5.10¹¹ vg) followed by a high-cholesterol diet for eight weeks (Figure 4A).

**Fig. 4.**
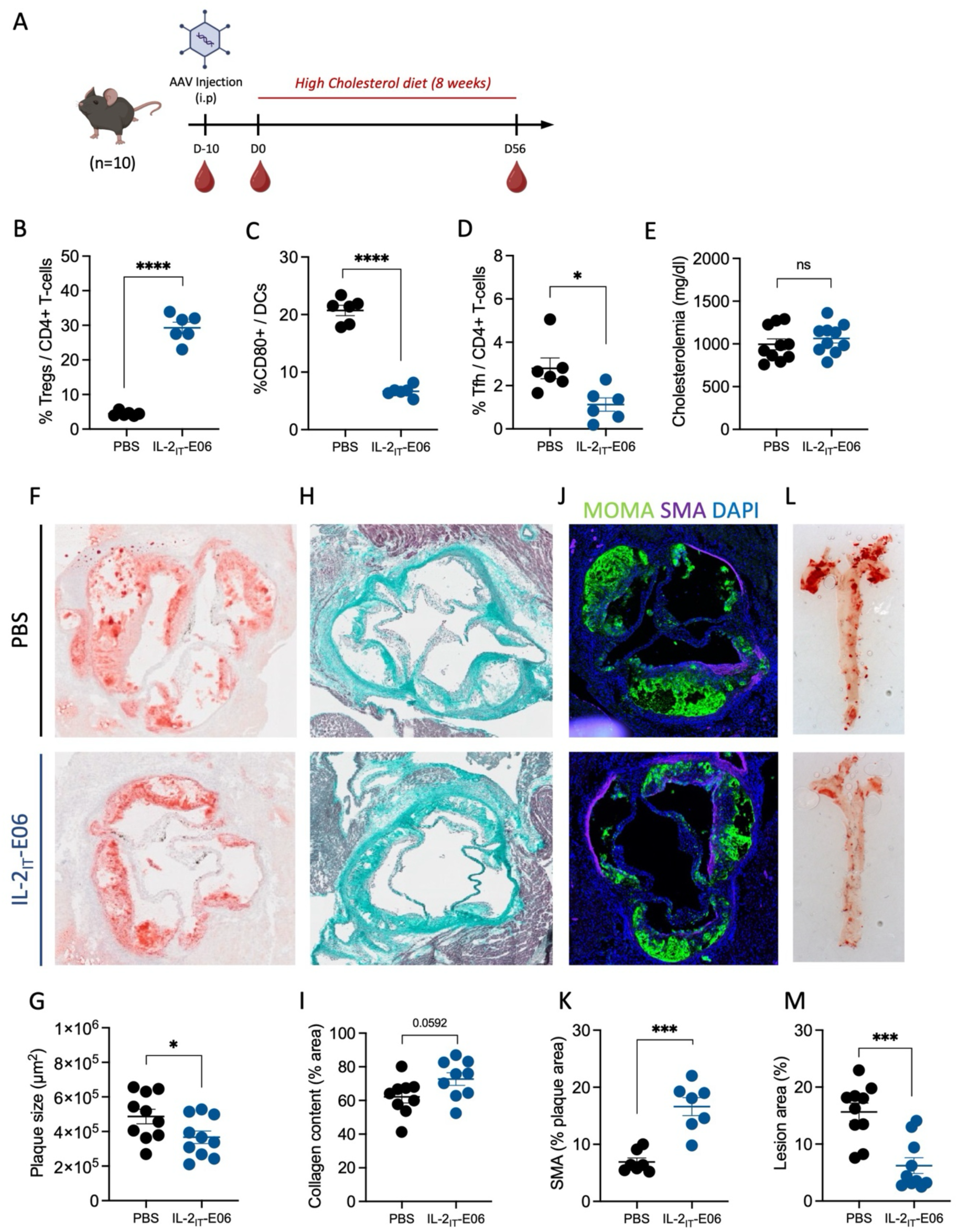
IL-2_IT_-E06 decreases lesions size and stabilizes atherosclerotic plaques. *Ldlr^-^*^/-^ mice (n=10/group) received i.p. administration of 5.10^11^ vg of either AAV-IL-2_IT_-E06 or PBS and kept on a chow diet for 10 days. After this time, mice received a high-cholesterol diet for 8 consecutive weeks to induce atherosclerosis **(A)**. Blood sampling was done before AAV injection, the day of diet change and after the 8 weeks of high-cholesterol diet. At the end of the experiment, mice were euthanized, aortas were collected and paraffin embedded, and spleen was collected to evaluate immune changes. Compared to untreated PBS group, IL-2_IT_-E06 increased Tregs **(B)**, decreased CD80 expression on DCs **(C)** as well as Tfh proportion **(D)** without any differences in cholesterolemia **(E)**. Aortas were cross-sectioned and atherosclerotic plaques were characterized to determine plaque size **(F,G)**, Collagen content **(H,I)**, SMA expression **(J,K)** to evaluate atherosclerosis severity. Longitudinal section of the whole aortas was also analyzed in *en face* preparations of the aortic arch and thoracic aorta to determine the full lesion areas **(L,M)**.

At the end of the experiment, the mice were euthanized, their aortas were collected, and atherosclerosis parameters were assessed using different staining procedures. Spleens were also processed to analyze immune parameters. IL-2_IT_-E06 expanded Tregs (5% vs. 30% among CD4⁺ T cells), reduced dendritic cells activation (20% vs. 5% of CD80+), and decreased T follicular helper cell (Tfh) frequency (2.5% vs. 1%, Figure 4B-D). Cholesterolemia remained comparable (≈ 1000 mg/dl, Figure 4E). IL-2_IT_-E06 significantly reduced lesion area (Figure 4F-G) and increased collagen (Figure 4H-I) and smooth muscle cell (α-SMA) content (Figure 4J-K)—hallmarks of plaque stabilization, while no difference was observed in macrophages composition (not shown). *En face* analysis of the aortic arch and thoracic aorta confirmed a significant reduction in lesion coverage (7% vs 17% in controls; Figure 4L-M). IL-2_IT_-E06 thus remarkably limits plaque burden and promotes stability. Plaque stabilization, rather than simple lesion reduction, represents a clinically meaningful endpoint for cardiovascular risk reduction.

### Affinity optimization enhances IL-2_IT_ potency

The binding affinity of the E06-scFv for OSE is much lower than that of IL-2 for the trimeric IL-2 receptors, potentially limiting the overall efficacy of the targeting. We aimed to increase the scFv affinity for OSE by replacing E06 with the higher-affinity M3C65 antibody, yielding IL-2_IT_-M3C65 (Figure 5A). Despite previous reports questioning the in vivo targeting efficiency of M3C65, we hypothesized that its higher affinity could be unmasked within the IL-2_IT_ format. Both IL-2_IT_-M3C65 and scFv-M3C65 bind specifically and equally well to their PC target (Figure 5B). SPR and NaSCN-ELISA showed that IL-2_IT_-M3C65 has a > 20-fold increased binding affinity for PC-BSA compared to IL-2_IT_-E06, with a Kd value of 1.8 nM and 37 nM respectively (Figure 5C-E). The modified construct bound PC-BSA but no other BSA-antigens (4-HNE-BSA, MDA-BSA, or BSA). It is also bound CuOx-LDL compared to native LDL (Figure 5F) and interestingly showed some cross-reactivity towards MAA-BSA, another class of OSE (Figure S3A). IL-2_IT_-M3C65 also stained both human and murine atherosclerotic plaques (Figure 5G; Figure S3B). Pharmacokinetics was comparable to IL-2_IT_-E06 (Figure 5H).

**Fig. 5.**
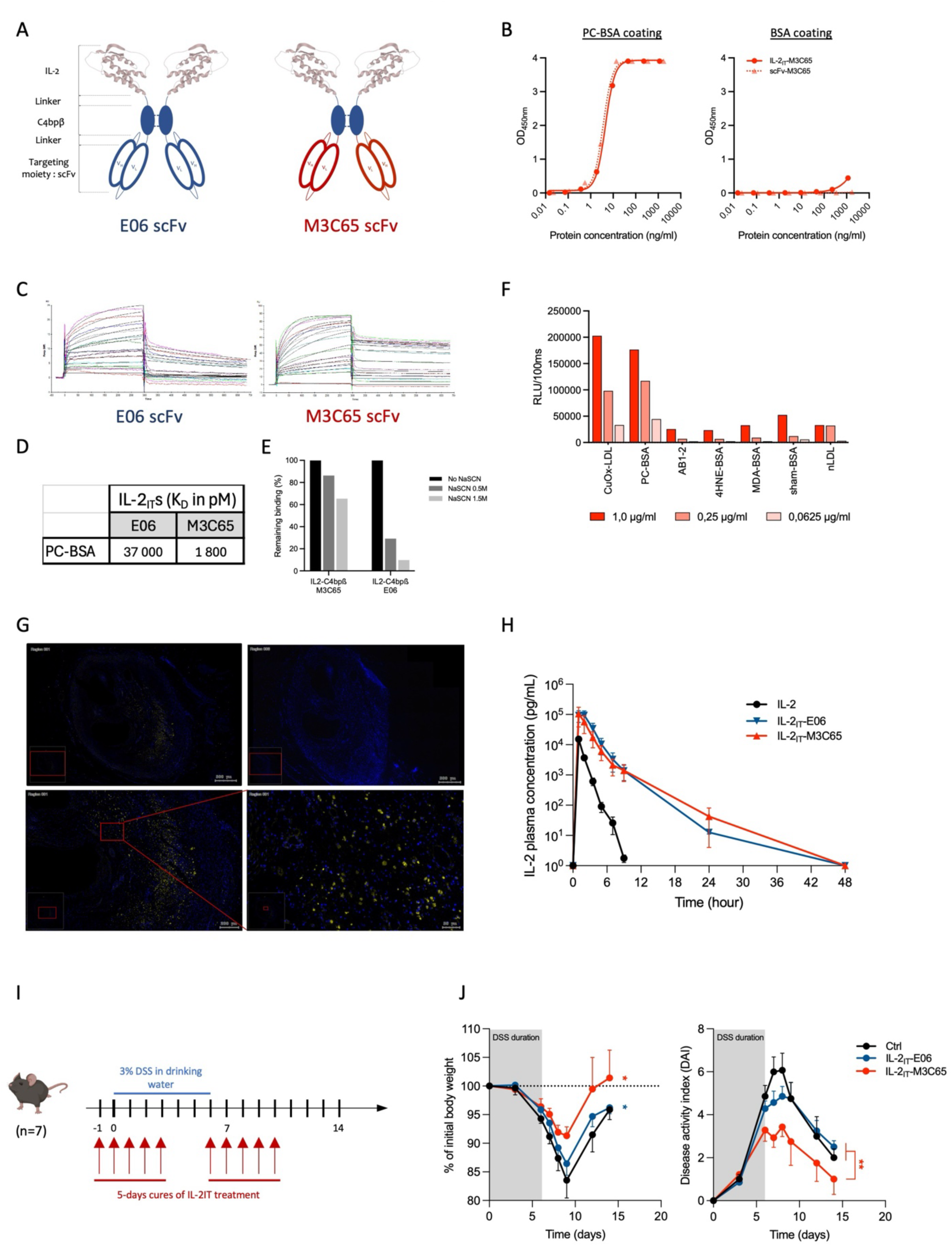
IL-2_IT_-M3C65: affinity optimization to enhance IL-2_IT_ potency. Schematic representation of IL-2_IT_-M3C65 after scFv replacement to enhance binding affinity **(A)**. IL-2_IT_-M3C65 and scFv-M3C65 bind specifically to PC-BSA and not to BSA alone in ELISA **(B)**. SPR kinetic for IL-2_IT_-E06 vs IL-2_IT_-M3C65 infused from 2nM to 200nM on PC-BSA **(C)**. KDs were computed and compared for both IL-2_IT_ proteins **(D)** where IL-2_IT_-M3C65 demonstrated increased affinity, confirmed by a NaSCN competitive ELISA binding experiment **(E)**. The binding specificity of IL-2_IT_-M3C65 was also evaluated against different OSEs (CuOx-LDL, idiotypic AB1-2, 4-HNE-BSA, MDA-BSA compared to sham-BSA and native LDL) at 1µg/ml, 0,25µg/ml and 0,0625µg/ml **(F)**. The binding capacity of IL-2_IT_-M3C65 was assessed in vitro on atherosclerotic cross-sections of human coronary arteries **(G)**. Control staining (upper right) were compared to full detection of IL-2_IT_-E06 staining (in yellow, upper left, lower left and right). Pharmacokinetics of IL-2_IT_-M3C65 was compared to IL-2_IT_-E06 and IL-2 after a single subcutaneous injection of 50.000IU *(H)*. C57BL/6J mice (n=7/group) received two cures of 5 daily injections of either IL-2, IL-2_IT_-E06 or IL-2_IT_-M3C65 proteins at 50.000 IU per injection subcutaneously **(I)**. 24 hours after the first injection, mice were given a 3% DSS solution as drinking water for 6 consecutive days. Colitis symptoms and severity were evaluated until D15 as bodyweight loss and DAI **(J)**.

In DSS-induced colitis, IL-2_IT_-M3C65, injected as protein, controlled weight loss and colitis severity, and is more efficient than IL-2_IT_-E06 (DAI score 3 vs. 4.5 respectively; Figure 5I-J).

Finally, as IL-2 therapy recently appeared to be particularly efficient to treat neuroinflammation, we aimed to assess IL-2_IT_-M3C65 in such a setting. In experimental cerebral malaria, a disease with a marked neuroinflammation, IL-2_IT_-M3C65 remarkably achieved 70% survival versus 40% for IL-2/dimIL-2 and 0% for controls, without negatively affecting parasitemia (Figure S4).

These results indicate that increasing scFv affinity enhances IL-2_IT_ efficacy across diverse inflammatory and infectious settings.

## Discussion

### First generation native IL-2: remarkable clinical achievements, yet with opportunities for optimization

Over the past decade, low-dose interleukin-2 has delivered a series of major clinical and translational advances. IL-2 is now firmly established as the first and only clinically validated means to safely and specifically stimulate regulatory T cells in humans, with reproducible biological activity and therapeutic benefit across diverse autoimmune, inflammatory, and neuroimmune diseases(*8, 15, 17, 39*). Although no phase 3 studies have yet demonstrated efficacy, these achievements have redefined IL-2 as a central immunoregulatory cytokine with broad therapeutic potential. Given these records, this has led to attempts at improving its overall performance.

### Second generation IL-2 muteins

For more than a decade, substantial efforts focused on re-engineering IL-2 to improve Treg selectivity(*40, 41*). Second generation IL-2 “muteins” were rationally designed or selected to reduce binding to IL-2Rβ (CD122), thereby favoring signaling through the high-affinity trimeric receptor (IL-2Rαβγ) enriched on Tregs(*20, 21, 42*). However, this approach came at a considerable cost: lowering IL-2Rβ affinity dramatically reduces IL-2’s intrinsic potency, making most IL-2 muteins being partial agonists. Although several IL-2 muteins expanded Tregs in humans, they consistently failed to produce meaningful clinical efficacy across autoimmune diseases. This dissociation between Treg expansion and therapeutic benefit likely reflects insufficient Treg activation rather than insufficient Treg numbers.

Of note, our dimIL-2 construct appears to have significantly improved properties compared to native IL-2 as well as to IL-2 muteins. As expected, it has a higher plasmatic half-life than IL-2, due to its increased size that abolishes urinary excretion. More unexpectedly, without any mutations in the IL-2 moiety, the dimerization process reduced the off-target activation of NK and Teffs. The particularity of our dimIL-2 may be explained by the two IL-2 moieties adopting a closed conformation that hides the IL-2Rβγ binding domain, as predicted by AlphaFold3 modelling. This interaction would likely reduce binding to the dimeric receptor while preserving binding to the trimeric IL-2R present on Tregs. These biological properties position dimIL-2 as an improved second-generation IL-2 compared to those in advanced clinical development.

Collectively, these experiences indicate that attenuating IL-2 to gain selectivity inherently weakens its therapeutic function, and that future IL-2–based therapies must avoid compromising the cytokine’s natural signaling strength. Improvement must therefore come not from weakening IL-2 biology, but from adding new functional properties while preserving wild-type potency.

### 3rd generation targeted IL-2

In oncology, the severe side effects of high dose IL-2 have led to the emergence of an alternative paradigm: immunocytokines, which fuse cytokines to antibodies that selectively bind to tumor-associated antigens or components of the tumor microenvironment(*43*). This strategy allows spatial precision—delivering potent cytokine signaling directly to diseased tissue while minimizing systemic toxicity. Several IL-2 immunocytokines have been developed(*44*), including constructs targeting tumor vasculature(*45*), extracellular matrix components(*46–48*), or tumor-associated integrins(*49*). These principles can be adapted to inflammatory diseases, which likewise exhibit characteristic molecular signatures at sites of chronic tissue damage.

We reasoned OSEs, widely exposed at sites of chronic inflammation, would constitute an ideal targeting signal for IL-2. OSEs accumulate on damaged membranes, apoptotic cells, oxidized lipoproteins, and extracellular matrix within inflamed tissue, creating highly specific molecular footprints of ongoing inflammation(*23, 24*). Anti-OSE antibodies such as the prototypic E06 have demonstrated anti-inflammatory properties, and previous studies—including work on atherosclerosis—showed that OSE targeting can efficiently localize scFvs to sites of pathology(*25, 37*).

Building on this rationale, we designed an Inflammation-Targeted IL-2 (IL-2_IT_) by fusing native IL-2 to a disulfide-linked dimeric IL-2 scaffold and an anti-OSE scFv. This construct preserves full IL-2 potency, exhibits prolonged plasma exposure due to increased molecular size, and selectively localizes to inflamed tissues in vivo. The IL-2_IT_ format thus represents a convergence of two principles: maintaining IL-2’s full biological activity and directing its action to pathological microenvironments where Treg activation is required. The IL-2_IT_ format therefore establishes a modular platform for inflammation-targeted cytokine delivery.

### Efficacy of IL-2_IT_ across models of autoimmunity and chronic inflammation

Across multiple disease models—psoriasis, colitis, atherosclerosis and neuroinflammation—IL-2_IT_ consistently outperformed native IL-2. In a model of psoriasis, IL-2_IT_ accumulated specifically at the site of imiquimod-induced inflammation and provided substantial clinical protection. In DSS colitis, IL-2_IT_ dramatically reduced weight loss and disease activity while selectively localizing within inflamed colonic tissue. Finally, in atherosclerosis, IL-2_IT_ not only reduced plaque burden but also enhanced plaque stability, a clinically relevant endpoint rarely achieved by immunotherapy(*50*). These observations indicate that targeted delivery of IL-2 to inflamed tissues simultaneously enhances efficacy and decreases the risk of exacerbating inflammation.

### Superior efficacy of IL-2_IT_-M3C65 despite previous concerns about targeting

To further optimize OSE targeting, we generated a high-affinity variant of Il-2_IT_ using the M3C65 anti-OSE antibody. Despite a prior report suggesting that M3C65 does not efficiently target inflammation in vivo(*51*), our IL-2_IT_ format revealed robust functional improvements upon incorporation of this scFv. IL-2_IT_-M3C65 displayed >20-fold higher affinity for OSEs, broader recognition of OSE classes, and preserved IL-2 pharmacodynamics. Functionally, IL-2_IT_-M3C65 demonstrated superior efficacy in DSS colitis, entirely preventing weight loss and markedly reducing disease activity. Moreover, in an experimental model of marked neuroinflammation—induced by cerebral malaria—IL-2_IT_-M3C65 increased survival to 70%, far surpassing IL-2 and controls, without altering parasite burden. These results highlight that affinity matters, as higher-avidity engagement of OSEs enhances therapeutic localization and efficacy. They position IL-2_IT_-M3C65 as a particularly potent next-generation IL-2 therapeutic.

### Perspectives

This work establishes inflammation-targeted IL-2 as bispecific Treg engager that represent a new class of “third-generation IL-2” therapeutics: from systemic immune modulation toward precision immunoregulation localized to sites of pathological inflammation. Unlike IL-2 muteins, which compromise potency to gain selectivity, IL-2_IT_ preserves full IL-2 biology while adding spatial precision and bifunctionality. The IL-2_IT_ platform appears versatile, mechanistically rational, and broadly applicable across autoimmune, inflammatory, cardiovascular, and neuroinflammatory disorders. IL-2_IT_ exemplifies a modular immunocytokine platform that can be tailored to disease-specific inflammatory signatures by exchanging scFvs or cytokines(*52*). This approach offers a promising pathway to unlock the full therapeutic potential of IL-2 in diseases with an inflammatory component.

## Materials and Methods

### Animals

Six to eight weeks old C57BL/6J female mice were purchased from Janvier Laboratory and maintained in our animal facility under specific pathogen-free conditions in agreement with current European legislation on animal care, housing, and scientific experimentation. The animals were housed in ventilated racks with an automatic watering system and ad libitum access to food. Temperature and humidity conditions are controlled and monitored. The lighting cycle is 12h per day (6AM-6PM). The animals are housed with their conspecifics (maximum of 5 per cage). At the end of the experiments, mice were euthanized by cervical dislocation.

For in vitro binding on murine atherosclerotic plaques, *Ldlr-/-* mice (MGI:2163421) were bred and kept in individually ventilated cages (IVC) with a 12-hour dark-/light-cycle and ad libitum access to sterilized food and water under barrier-specific pathogen-free conditions (SPF) at the Core Facility for Animal Breeding and Husbandry of the Center for Biomedical Research at the Medical University of Vienna, Austria. To induce atherosclerosis, ten-week-old female mice were placed on an atherogenic Western-type diet containing 21% milk fat and 0.21% cholesterol (TD88137, Ssniff Spezialdiäten GmbH) with ad libitum access to sterilized pellets for 12 weeks. Mice were euthanized and aortic root sections were prepared as previously described(*53*). All experimental studies were approved by the Animal Ethics Committee of the Medical University of Vienna and the Austrian Federal Ministry of Education, Science and Research, and were performed according to Good Scientific Practice and national and international institutional guidelines (License number BMWF 2021-0.030.759).

For atherosclerosis experiments, eight-week-old female Ldlr−/− mice (purchased from Jax/Charles River) were injected intraperitoneally with 5.10¹¹ vg of AAV-IL-2_IT_-M3C65 or PBS. Ten days after injection, the mice were placed on a high-cholesterol diet (HCD) containing 15% fat, 1.25% cholesterol and 0% cholate.

For the NHP experiment, four adults Mauritian cynomolgus macaques (*Macaca fascicularis,* two males and two females) was included after being detected seronegative for AAV. Macaques will be housed in groups within IDMIT animal facilities at CEA, Fontenay-aux-Roses. Water and food were provided to the animals ad libitum. All experimental procedures were conducted according to European guidelines for animal care and use for scientific purposes (Directive 63-2010, “Journal Officiel des Communautés Européennes”, L276, September 22, 2010). The study was approved by the local ethical committee under the number A20_037 and by the French Administration (APAFIS#24434-2020030216532863 v3).

### Fusion proteins

IL-2 fusion proteins were produced by Icosagen with its proprietary QMCF Technology platform. Proteins were engineered with a C-terminal Twin-Strep-tag or 6xHisTag for affinity purification, and their coding sequences were cloned into pQMCF expression vectors. Sequence-verified vectors were chemically transfected into CHOEBNALT-1E9 cells using reagent R007, and cells were cultivated in CHO TF Medium (Sartorius) supplemented with HyClone™ ActiPro™ cell culture media and HyClone™ Cell Boost™ 7a and 7b feeds (Cytiva) as per Icosagen QMCF Technology protocol. Following production, the secreted proteins were purified from cell culture supernatant via affinity chromatography, using StrepTactinXT 4Flow Cartridge columns (IBA Lifesciences). Final polishing was done by preparative size exclusion chromatography on Superdex 200 columns (Cytiva), and proteins were formulated into PBS pH 7.4 buffer. Proteins were then resuspended in PBS or Citrate buffer when appropriate, aliquoted and concentrated at 1.5-2.5 mg/ml and stored at 4°C. All the proteins were administered in mice by subcutaneous injections in 100µl.

### rAAV generation and *in vivo* administration

AAV vectors were generated by the AAV Vector Unit at ICGEB Trieste (http://www.icgeb.org/RESEARCH/TS/COREFACILITIES/AVU.htm). Briefly, HEK293 cells were used to produce AAV8 particles expressing IL-2_IT_-E06, IL-2_IT_-M3C65, using the AAV Helper Free Packaging System (Cellbiolabs, #VPK-402). The therapeutic gene was cloned and expressed under the control of cytomegalovirus (CMV) immediate-early enhancer and promoter. After production, viral stocks were purified through CsCl2 gradient centrifugation. The titration of the viral stocks was determined by quantifying the number of viral genomes per ml, by real-time PCR, as described in(*54*).

### Protein modelling

The structure of the dimIL-2 complex was predicted using AlphaFold 3 (AF3)(*55*). Default AlphaFold 3 parameters were used unless otherwise specified. Predicted models were ranked according to the AlphaFold confidence metrics, including the predicted local distance difference test (pLDDT) and inter-chain predicted alignment error (PAE). The top-ranked model was selected for further analysis.

Structural visualization and analysis were performed using PyMOL (The PyMOL Molecular Graphics System, Version 3.10 Schrödinger, LLC). Molecular surfaces were generated to highlight exposed regions and interaction sites. Conserved IL-2 residues known to be involved in IL-2 receptor α and β chain interactions were mapped onto the predicted structure for comparative analysis. Disulfide bond formation between C4BPβ subunits was assessed based on cysteine proximity and geometry in the predicted model. Key residues of interest were highlighted using stick and color representations to facilitate interpretation of functional sites and dimerization interfaces.

Protein–protein interactions within the predicted complex were analyzed using the Protein–Ligand Interaction Profiler (PLIP)(*56*). The predicted structure was provided as input to PLIP to identify non-covalent interactions, including hydrogen bonds, hydrophobic contacts, salt bridges, and π-related interactions. Interaction networks were extracted and visualized to support qualitative assessment of interface regions. All interactions reported are derived from the predicted structural model and should be interpreted as computationally inferred interactions rather than experimentally validated contacts.

IL-2_IT_-E06 modelling was performed using MOE modelling tool to assemble the three components together (IL-2, C4bpβ and E06). IL-2 alone and in complex with the three subunits of IL-2 receptor was obtained from PDB 2B5I, dimerization of C4bpβ from AlphaFold F3 and E06 from MOE. Closed IL2/IL2 conformation was obtained from PDB ID 1QVN.

### pSTAT5 assay

Human blood samples from healthy volunteers were obtained from Etablissement Français du Sang (EFS) in Paris, France. Informed consent was obtained from each volunteer.

The effects of human IL-2, fusion proteins and targeted proteins on the induction of STAT5 phosphorylation (pSTAT5) were assessed in human CD4+ regulatory T cells (Treg; CD4+Foxp3+CD127lo/-), CD4+ conventional T cells (Tconv; CD4+Foxp3-), CD8+ T cells and natural killer cells (CD3-CD56+) using flow cytometry. Ten-fold dilution of human IL-2, fusion and targeted proteins mixed with 100µl of whole blood for 15min at 37°C were performed before pSTAT5 staining using the Phospho-Epitopes exposure kit (PERFIX EXPOSE kit, Beckman Coulter). In conjunction with the intracellular pSTAT5 signal, surface markers allowed the comparison of activity and the calculation of half maximal effective concentration (EC_50_) values for different lymphocyte cell populations.

### ELISA

To measure plasmatic and urinary concentrations of the different dimIL-2 and IL-2_IT_s, blood samples were collected in heparin tube (mice samples). Plasma was isolated by centrifugation (800rcf; 10 minutes), frozen and kept at −20°C until use. Level of plasmatic and urinary IL-2 were measured according to the manufacturer’s recommendations using anti-human IL-2 ELISA kit (Thermofisher). Briefly, 100 µL of IL-2 capture antibody (1:250; 100µL/well) was coated in flat-bottom, 96-well plates and incubated overnight at 4°C. After washing and blocking non-specific binding sites with PBS + Tween (0.05%) + 1% bovine serum albumin (BSA) for 1 hour at room temperature (RT), 100µL of diluted samples performing 5-fold serial were added to each well and incubated for 2 hours at RT. Then, plates were incubated one hour with detection antibody (1:250; 100µL/well) before revelation using streptavidin-HRP concentrate (1:250; 100µL/well). For IL-2_IT_s, we used capture anti-IL-2 monoclonal antibody (MQ1-17h12) or Phosphocholine-BSA (LGC Biosearch technologies) coated at 1µg/mL and detection performed using anti-His antibody (1:1000; Thermofisher) at 1µg/mL before revelation with ultrasensitive streptavidin-HRP (1:2000; Sigma Aldrich).After adding TMB solution (RT; 15 min) and stop solution, the plate was read at 450nm using an automatic ELISA plate reader (DTX 880, Multimode detector; Beckman Coulter).

For OSE binding screening, 96-well plates were coated with 50 µl/well of antigens diluted in PBS containing 0.27 mM EDTA (pH 7.4; GIBCO 14190-094), including PC-BSA (5 µg/ml, PC-1011H-10-BS), MDA-LDL (5 µg/ml, self-prepared), CuOx-LDL (5 µg/ml, self-prepared), AB1-2 (anti-idiotypic T15/E06 antibody; 2 µg/ml, BIOTEM), MAA-BSA (5 µg/ml, self-prepared), MDA-BSA (5 µg/ml, self-prepared), 4HNE-BSA (5 µg/ml, self-prepared), sham-treated BSA (5 µg/ml, self-prepared), and native LDL (5 µg/ml, isolated from healthy donors). Plates were incubated overnight at 4°C, for 60 min at 37°C, or for 120 min at room temperature (RT) on a shaker. After coating, plates were washed with PBS/EDTA (Bio-Tek plate washer) and blocked with 100 µl/well of 1% BSA in TBS/EDTA (pH 7.4, 0.27 mM EDTA) for 60 min at RT or overnight at 4°C. Following another wash, M3C65-IL2_IT_ and E06-IL2_IT_ were diluted to the indicated concentrations in 1% BSA-TBS/EDTA were added in triplicate (3 × 25 µl/well), and plates were incubated overnight at 4°C or for 120 min at RT on a shaker. After washing, 50 µl/well of secondary antibody (Strep-Tactin AP, 1:2000 dilution; IBA-2-1503-001) in 1% BSA-TBS/EDTA was added and incubated for 120 min at RT or overnight at 4°C. Plates were then washed with TBS (pH 7.4), followed by addition of 25 µl/well of Lumiphos Plus (33%, Lumigen P701, diluted 1:2 in sterile diH₂O). After 60 min incubation at RT in the dark, chemiluminescence was measured (RLU/100 ms) using a BIOTEC H1 ELISA Reader (Gen5 software v1.31).

### Histology

Immunohistochemistry was performed to assess the binding of IL-2_IT_-E06 and IL-2_IT_-M3C65 to atherosclerotic lesions. After dewaxing and rehydration, human and murine slides were subjected to antigen-retrieval for 20-30 minutes at 97,5-99°C using Citrate Antigen Retrieval Solution (Sigma-Aldrich) and PBS-T washing. Cross sections were blocked at room temperature using 2% BSA-PBS-Tween20 for 1 hour. Sections were incubated with 4µg/ml M3C65-IL2_IT_ or 8µg/ml E06-IL2_IT_ in antibody dilution buffer (containing 2% BSA in 0.01M PBS, PH7.2) at 4°C overnight followed by incubation with DY-649-conjugated Strep-TactinXT (IBA-2-1568-050, IBA Life Sciences) diluted 1:200 in PBS with 1-2% BSA. Slides were mounted with Fluoromount before image acquisition on a TFSpectra TissueFAXS (TissueGnostics) using the TissueFAXS Viewer 7.1 Software. Selected sections were acquired on an LSM900 Confocal Microscope (Carl Zeiss AG) using the Zen3.4 Pro Software.

Aortic roots were embedded in OCT and frozen immediately for subsequent sectioning and staining. Hearts were sectioned through the aortic root and stained with Oil Red O (Sigma-Aldrich) for quantification of lesion area. Collagen content and necrotic core surface was quantified after Masson’s Trichrome staining.

For immunostaining, slides were fixed with 4% formaldehyde, permeabilized with 0.1% Triton X-100, and blocked with 5% BSA-TBS for 1hour at RT. The sections were incubated with primary antibodies against MOMA (MAB1852, Millipore) and SMA (F3777, Sigma-Aldrich). Sections were then incubated with appropriate secondary antibodies and stained with DAPI to detect nuclei. The images were taken with Olympus Slide View VS200 and analyzed using QuPath software.

The aortas were dissected under a microscope and fixed in 4% PFA, opened, stained with Oil Red O, flattened and images of the aortas were captured and quantified by analysis of the entire *en face* aorta as previously described(*57*).

### Experimental autoimmune encephalomyelitis

C57BL/6J mice were injected subcutaneously in the posterior right and left flank with an emulsion containing 200µg of MOG_35-55_ supplemented with 500µg of *Mycobacterium tuberculosis* (BD biosciences) in complete Freund adjuvant (CFA) (Sigma-Aldrich). On the day of immunization and 2 days after, mice received 400ng of Pertussis toxin (Enzo laboratory) by intraperitoneal injection. Mice were scored daily for clinical signs of EAE beginning on day 7 after injection as follow: 0, no clinical expression of disease, 1, limp tail without hind-limb weakness; 2, limp tail and weakness of hind legs with movement difficulties; 3, limp tail and paralysis of hind legs and one front leg; 3.5, limp tail and paralysis of hind legs and front legs; 4, moribund.

### DSS-induced Colitis model

DSS-induced colitis was induced in C57BL/6J mice by oral administration of 3% of Dextran Sulfate Sodium (DSS 40,000Mw; Sigma-Aldrich) in drinking water for 6 days. Mice were monitored daily for body weight, consistency and presence of blood in their stool until day 15. The following scoring system has been used to evaluate the severity: Loss of body weight between 0-5%: 1, 5-10%: 2, 10-15%: 3 and over 15%: 4. Stool consistency: normal stool: 0; formed but soft stool: 1; loose stool: 2; mild diarrhea: 3; watery diarrhea: 4. Fecal bleeding: absence: 0, presence: 2, gross bleeding: 4.

### Psoriasis model

C57BL/6J mice was shaved in the back prior to psoriasis induction. Mice received 62,5 mg of 5% imiquimod cream on the shaved back as well as a small quantity on one ear once a day for 6 days. The second ear received Vaseline as control. Skin thickness on the back and the ears was measured every day using an electronic caliper, and evolution was reported as % of baseline. Scaling was also measured as well as redness on the back to determine skin inflammation and lesion (score 0 to 4).

### Atherosclerosis

Atherosclerosis was induced in eight-week-old female Ldlr−/− mice injected intraperitoneally with 5.10¹¹ vg of AAV-IL-2_IT_-E06 or PBS. Ten days after injection, the mice were placed on a high-cholesterol diet (HCD) containing 15% fat, 1.25% cholesterol and 0% cholate for 8 consecutive weeks. At the end of the experiment, Blood was obtained through retro-orbital bleeding. Plasma was separated by centrifugation at 13,000 rpm and kept at −20 °C. Cholesterol levels were measured using DiaSys^®^ Cholesterol FS* kit (DiaSys Diagnostics). Spleen were harvested to determine pharmacodynamics, whereas aortic roots and aortas were dissected to evaluate atherosclerosis severity.

### Statistical analyses

Graphing and statistical analyses were performed using GraphPad Prism 10.5.0 software. Statistical analyses were performed using the unpaired two-tailed Mann-Whitney test, Kruskal-Wallis test or a one-way ANOVA followed by a Tukey’s multiple comparison test when appropriate. When two groups only were compared, a Shapiro-Smirnov test was used to determine Normality. An Unpaired t-test or a Welch’s t-test was then used when appropriate depending on the result obtained with a F-test. Data were expressed as mean +/- SEM. Statistical significance was taken at the 5% level (p < 0.05), and the degree of significance was indicated as follows: *p<0.05; **p<0.01; ***p<0.001, ****p<0.0001. Only significant differences are shown unless non-significant differences (denoted as “ns”) are specifically indicated.

## List of supplementary materials

Supplementary materials and methods

Fig. S1 to S4

## Acknowledgement

We thank all the partners of the TiilT consortium (European Union’s Horizon Europe Program N°101072435) who participated to this work. We thank N. Bartoloni and I. Del Giudice for AAV production, Y. Guan for help in murine Colitis experiments, C. Aheng for managing the TiilT project and the CEF – UMS28 for mice management. We also thank M. Giacca, M. Tomczyk, J. Humrich, R. Akbarzadeh and T. Brodiazhenko, all TiilT partner, for discussions. Part of the figures were created with BioRender.com

## Authors contribution

Conceptualization: DK

Funding acquisition: DK

Investigation: TV, MCM, NB, FP, MS, FS, CA, LZ, TB, LP, JT, IL

Formal analysis: CA, JT

Supervision: DK, SZ, RLG, MR, ZM, CB

Writing – original draft: TV, DK

Writing – review & editing: all authors

