## Supplementary methods and figures for "Targeting IL-2 to inflamed tissues via oxidation-specific epitopes enables third-generation bispecific IL-2 therapeutics"

Thomas Vazquez *et al.*

### Experimental Cerebral Malaria

**C57BL6/6 mice was treated with $10^{11}$ vg AAV-IL-2, AAV-dimIL-2 or AAV-IL-$2_{IT}$-M3C65 two weeks prior to *Plasmodium berghei* ANKA parasites i.p. infection.** Mortality was followed from ECM window (6.5 to 10 days post-infection) and clinical signs of ECM were registered.

### Flow cytometry and antibodies

After mice blood hemolysis, isolated immune cells were stained with the following antibodies at predetermined optimal dilutions for 20 minutes at 4°C: CD3, CD4, CD8, CD25, NKp46, and CD19. Intracellular detection of Foxp3 was performed on fixed and permeabilized cells using FoxP3 staining buffer kit (eBioscience FoxP3/Transcription). Cells were acquired on cytoflex LX (Beckman Coulter) and analyzed with FlowJo software. Doublets were excluded by forward/side scatter gating. CD4+ Tregs were defined as CD4+CD25+Foxp3+ cells, CD4+ Teffs as CD4+CD25+Foxp3- cells also called Tconv CD4+CD25+, Natural killer cells as Nkp46+ cells and Bcells as CD19+ cells.

For pSTAT5 assay, human whole blood was incubated with the following antibodies against membrane proteins for 15min: CD3, CD127, CD4 and CD8 and CD56-APC. After fixation and permeabilization, cells were stained with intracellular p-STAT5 and Foxp3 antibodies.

### pSTAT5 assay

Human blood samples from healthy volunteers were obtained from Etablissement Français du Sang (EFS) in Paris, France. Informed consent was obtained from each volunteer.

The effects of human IL-2, fusion proteins and targeted proteins on the induction of STAT5 phosphorylation (pSTAT5) were assessed in human CD4+ regulatory T cells (Treg; CD4+Foxp3+CD127lo/-), CD4+ conventional T cells (Tconv; CD4+Foxp3-), CD8+ T cells and natural killer cells (CD3-CD56+) using flow cytometry. Ten-fold dilution of human IL-2, fusion and targeted proteins mixed with 100µl of whole blood for 15min at 37°C were performed before pSTAT5 staining using the Phospho-Epitopes exposure kit (PERFIX EXPOSE kit, Beckman Coulter). In conjunction with the intracellular pSTAT5 signal, surface markers allowed the comparison of activity and the calculation of half maximal effective concentration ($EC_{50}$) values for different lymphocyte cell populations.

**Ethical approval**

Human leukapheresis products were obtained from the Etablissement Français du Sang (EFS) under the agreement n°21/EFS/024 established between EFS and our institution. Written informed consent was obtained from all healthy volunteers by EFS in accordance with ethical guidelines.

For murine experiment, the animal procedures were approved by the "Comité d'éthique en expérimentation animale Charles Darwin" registered at the "Comité National de Réflexion Ethique sur l'Expérimentation animale", CEEAACD 005, under the project agreement n°29481. For in vitro binding on murine atherosclerotic plaques, all experimental studies were approved by the Animal Ethics Committee of the Medical University of Vienna and the Austrian Federal Ministry of Education, Science and Research, and were performed according to Good Scientific Practice and national and international institutional guidelines (License number BMWF 2021-0.030.759).

For NHP experiment, all experimental procedures were conducted according to European guidelines for animal care and use for scientific purposes (Directive 63-2010, "Journal Officiel des Communautés Européennes", L276, September 22, 2010). The study was approved by the local ethical committee under the number A20_037 and by the French Administration (APAFIS#24434-2020030216532863 v3).

### Supplementary figures and legends

A

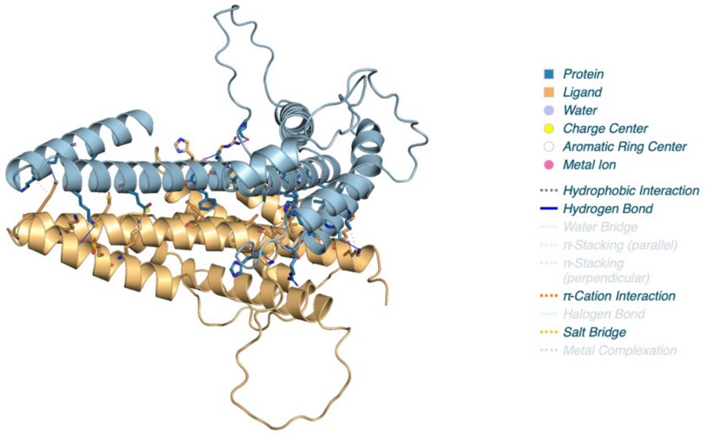


B

Hydrophobic interactions

| Index | Residue | AA | Distance |
|---|---|---|---|
| 1 | 8B | LYS | 3.58 |
| 2 | 12B | LEU | 3.73 |
| 3 | 12B | LEU | 3.12 |
| 4 | 19B | LEU | 3.49 |
| 5 | 85B | LEU | 3.72 |
| 6 | 85B | LEU | 3.80 |
| 7 | 88B | ASN | 3.94 |
| 8 | 91B | VAL | 3.79 |
| 9 | 92B | ILE | 3.26 |
| 10 | 163B | ALA | 3.99 |
| 11 | 164B | PHE | 3.86 |
| 12 | 164B | PHE | 3.01 |
| 13 | 182B | LYS | 3.97 |
| 14 | 192B | LYS | 3.72 |

Hydrogen bonds

| Index | Residue | AA | Distance H-A |
|---|---|---|---|
| 1 | 8B | LYS | 1.80 |
| 2 | 15B | GLU | 3.12 |
| 3 | 79B | HIS | 2.13 |
| 4 | 81B | ARG | 2.13 |
| 5 | 87B | SER | 2.01 |
| 6 | 88B | ASN | 2.54 |
| 7 | 153B | PRO | 3.37 |
| 8 | 156B | GLU | 1.97 |
| 9 | 159B | LYS | 1.84 |
| 10 | 182B | LYS | 2.95 |
| 11 | 192B | LYS | 2.73 |

Salt bridges

| Index | Residue | AA | Distance |
|---|---|---|---|
| 1 | 9B | LYS | 3.12 |
| 2 | 95B | GLU | 3.44 |
| 3 | 175B | GLU | 3.01 |
| 4 | 182B | LYS | 4.38 |
| 5 | 196B | GLU | 4.56 |
| 6 | 199B | LYS | 2.86 |

Π -Cations interactions

| Index | Residue | AA | Distance |
|---|---|---|---|
| 1 | 16B | HIS | 1.23 |

C

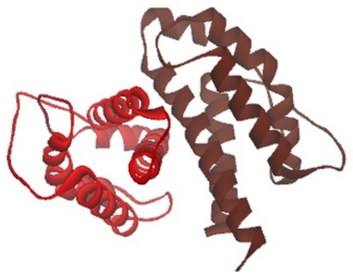

D

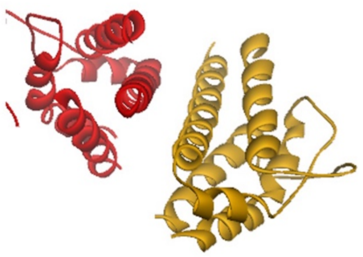

**Fig. S1. dimIL-2 and IL-2 homodimerization modelling.** IL-2/IL-2 predicted conformation from dimIL-2 is shown **(A)** with a description of the interactions involved in the IL-2 dimerization. Tables illustrating Hydrophobic interactions, Hydrogen bonds, Salt bridges and Π-Cations interactions are shown **(B)** with the residues and their positions, as well as their distance.

Conformation of IL-2 homodimer from AlphaFold 3 prediction **(C)** and the crystal structure from PDB ID 1QVN **(D)** are shown for comparison.

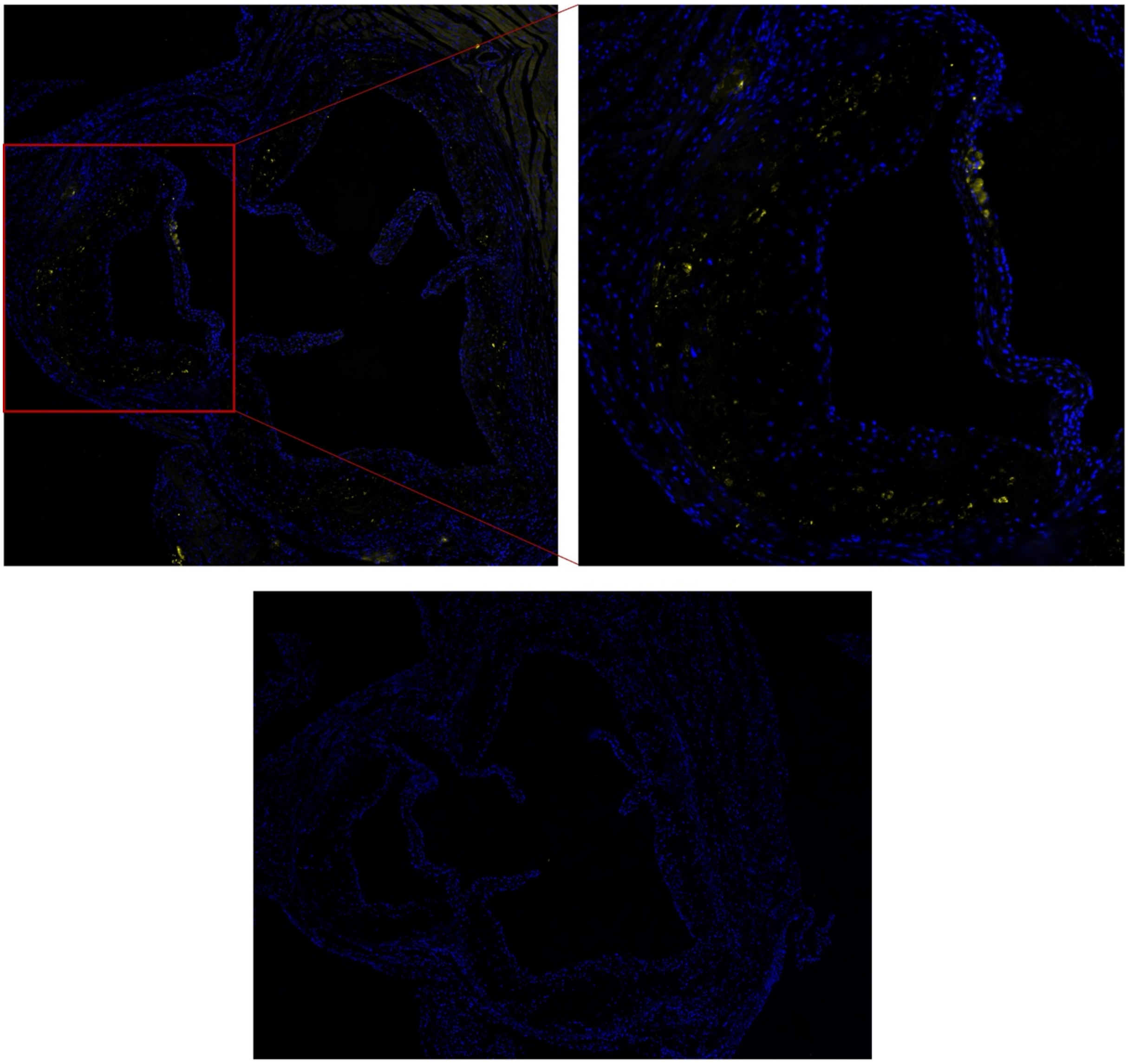

**Fig. S2. IL-2$_{IT}$-E06 binds to murine atherosclerotic plaques.** Murine atherosclerotic plaques from *Ldlr-/-* mice induced by high-cholesterol diet was paraffin-embedded and cross-sectioned. Staining using StreptactinXT (yellow, upper left and right) or control (bottom) were performed to detect OSE binding in atherosclerotic plaques.

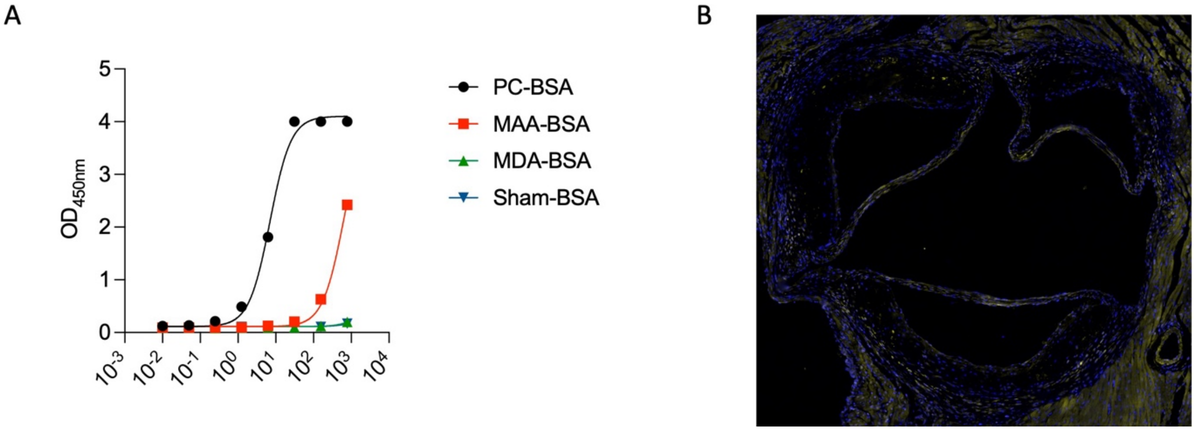


**Fig. S3. Binding properties of IL-2$_{IT}$-M3C65.** A dose-response of IL-2$_{IT}$-M3C65 protein was evaluated for OSE binding against different OSE-BSA proteins **(A)** : PC-BSA (black), MAA-BSA (red), MDA-BSA (green) or sham-BSA (blue). Murine atherosclerotic plaques induced by high-cholesterol diet feeding of *Ldlr-/-* mice was paraffin-embedded and cross-sectioned. Staining using StreptactinXT (yellow) **(B)** were performed to detect OSE binding in atherosclerotic plaques.

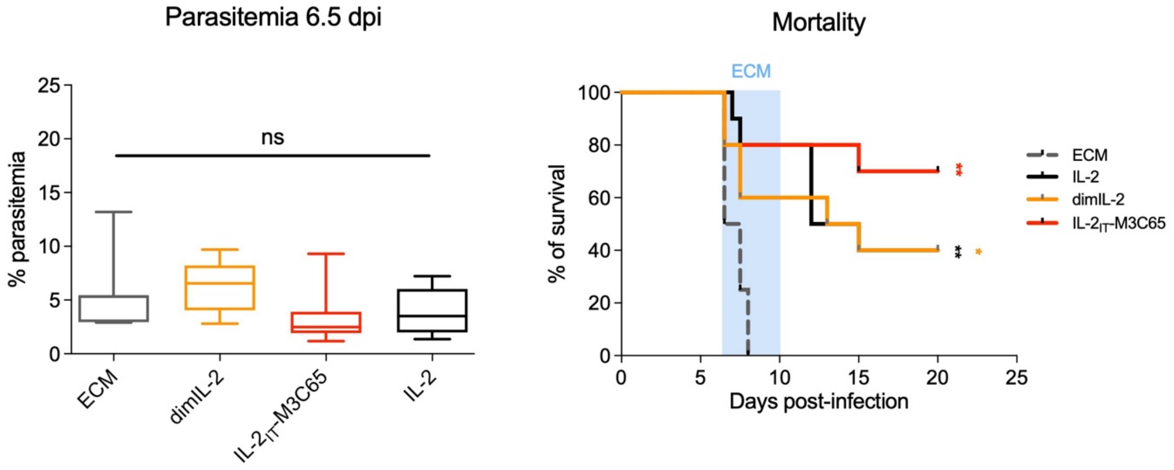


**Figure S4. Efficacy of IL-2$_{IT}$-M3C65 in Experimental Cerebral Malaria. C57BL6/6J (n=10/group) mice were treated with $10^{11}$ vg AAV-IL-2, AAV-dimIL-2 or AAV-IL-2$_{IT}$-M3C65 two weeks prior to *Plasmodium berghei* ANKA parasites ip infection.** Parasitemia at 6.5 dpi (left) and mortality (left) was followed from ECM window (6.5 to 10 days post-infection) and clinical signs of ECM were registered.
